# An updated assessment of the genomic health of *Odocoileus* [deer]

**DOI:** 10.64898/2026.07.13.738275

**Authors:** B Cars, A.B.A Shafer

## Abstract

Genomic health estimates help inform conservation and management decisions, with genetic load and runs-of-homozygosity (ROH) being two key metrics. White-tailed deer (*Odocoileus virginianus*) and mule deer (*O. hemionus*) are found throughout North America, with some populations declining or of conversation concern. Using genome-wide data from samples across their range, we provide the first estimate of genetic load in mule deer, and revisit ROH estimates using model-based approaches. These updated estimates of ROH notably showed elevated inbreeding in the Key deer, consistent with current conservation designations. We also detected a relatively high number loss-of function mutations in mule deer that we attributed to historical bottlenecks. We also observed an increased overall genetic load in *O. hemionus* from the Pacific Northwest.

## Introduction

Genome-scale data can provide important insights into the health of organisms. Metrics like genetic load and runs of homozygosity (ROH) have garnered interest in populations managed for conservation or harvest (Schmidt et al. 2024; Kyriazis et al. 2025), as there are documented links to fitness (Stoffel et al. 2021) and population viability (Kardos et al. 2023). With any new data, methods evolve, and two distinct approaches – rule and model-based – for estimating ROH have emerged (see Shafer & Kardos 2025). Importantly, rule-based approaches can be biased with moderate to low coverage data (e.g. Shi et al. 2026; Taylor et al. 2026), leading to the recommendation to use model-based ROH estimates in free-ranging populations (Shafer & Kardos 2025). And while genetic load predictions were first developed in humans (Henn et al. 2015), they have improved and crossed over to non-model organisms (Bertorelle et al. 2022). Of relevance is that load is generally positively associated with population size (van der Valk et al. 2019), but demographic effects can have an impact (Wootton et al. 2023), with purging appearing to be relatively common.

White-tailed deer (WTD; *Odocoileus virginianus*) and black tailed / mule deer (MD; *O. hemionus*) are widespread ungulates in North America, though WTD were severely reduced, and extirpated in some regions, at the turn of the 20^th^ Century, but have since recovered (Webb et al. 2018). Mule deer now experiencing declines throughout their range (Bergman et al. 2015). Both species have had genome-wide inbreeding assessed using rule-based ROH metrics (Kessler et al. 2024), with the endangered Key deer not showing elevated levels (Cars et al. 2024). WTD show high genetic load consistent with predictions (Wootton et al. 2023), but no estimates have been provided for mule deer, which are of management concern (Bergman et al. 2015). Here we provided updated model-based estimates of inbreeding in both species, and the first estimate of genetic load in MD.

## Methods

### Sampling

Genome sequence data from 19 *Odocoileus hemionus* individuals, including both mule deer and black-tailed deer, were included in this study (PRJNA830519). Mule deer and black-tailed deer were grouped together as we calculated individual metrics and sample size is small for the latter (but see Kessler et al. 2024 for population metrics). Available genome sequence data from 51 previously published white-tailed deer (*Odocoileus* virginianus) genomes were also included for comparative analyses (Cars et al., 2024). The white-tailed deer samples included 16 samples from Saint-Pierre and Miquelon (France, but coastal Canada), 21 from the North American mainland, 10 from the Florida Keys (USA) and 4 from Anticosti Island (Quebec, Canada; Figure 1).

**Figure 1.**
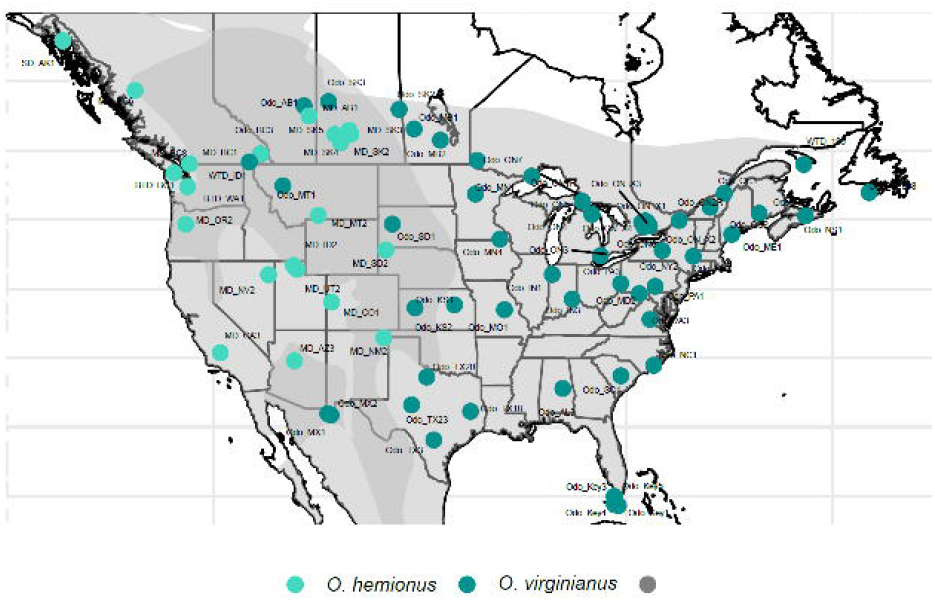
Range wide sampling of White-tailed deer (*Odocoileus virginianus*) and mule deer (*O. hemionus*) used in this study.

### Read alignment and variant calling

Raw sequencing reads were first assessed for quality using FastQC (v0.11.9; Andrews, 2010). Low-quality bases were removed with Trimmomatic (v0.39; Bolger *et al*., 2014), after which the filtered reads were aligned using BWA-MEM (v0.7.17; Li, 2013) to the appropriate annotated reference genome (*O. hemionus*, GCA_020976825.1), or *O. virginianus*, GCA_023699985). Resulting alignments were sorted using SAMtools sort (v1.15.1; Li *et al*., 2009). Duplicate reads were identified with Picard MarkDuplicates (v2.23.2; Broad Institute, 2019) and removed using Sambamba (v0.8.2; Tarasov *et al*., 2015). To verify sequencing depth and genome coverage, alignment metrics were summarized using SAMtools coverage and depth, as well as Sambamba flagstat. Additionally, depth and breadth were calculated with mosdepth (v0.3.1; Pedersen and Quinlan, 2018).

Single-nucleotide polymorphisms (SNPs) were identified using ANGSD (v0.939; Korneliussen *et al*., 2014). Genotype likelihoods were estimated under the GATK model framework (gl 2) and output in BEAGLE format (doGlf 2). Sites were retained as SNPs only if they met filtering criteria, including a minimum significance threshold of p□<□1□×□10□□ (-SNP_pval 1e-6), a minimum mapping quality of 20 (-minMapQ 20), and a minimum base quality of 20 (-minQ 20). Genotypes were then generated for retained sites (doGeno 4). Major and minor alleles were inferred directly from genotype likelihoods (doMajorMinor 1), with allele frequencies assumed to be known (domaf 1).

### Subpopulation Diversity, ROHs, and SNP Effects

We estimated population diversity summary statistics following the same approach as described in Cars et al. (2024), ensuring that values were directly comparable across studies. Nucleotide diversity (π) was calculated for each subpopulation using VCFtools (v0.1.16; Danecek *et al*., 2011) in 10□kb windows (-window-pi 10000). Tajima’s D was estimated for each subpopulation, again using 10□kb windows (-TajimaD 10000) and considering only bi-allelic sites. Genetic differentiation between populations was quantified using fixation indices (FST), calculated once again in 10□kb windows (-fst-window-size 10000).

Runs of homozygosity (ROHs) were identified for all individuals (MD and WTD) using bcftools roh (v1.22; Narasimhan *et al*., 2016; Danecek *et al*., 2021). ROHs in the white-tailed deer dataset were previously estimated using PLINK (Cars *et al*., 2024). ROH segments were filtered to include only regions with a minimum length of 100□kb, at least 50 SNPs, and a quality score ≥□25. For each individual, total ROH length and the number of ROH segments were calculated from filtered outputs. Mean ROH length was obtained by dividing total ROH length by the number of segments. Genomic inbreeding coefficients (FROH) were estimated as the proportion of the genome contained within ROHs, calculated from total ROH length divided by the autosomal genome size.

We assessed genetic load using SnpEff (Cingolani *et al*., 2012) by quantifying homozygous missense mutations, loss-of-function (LOF) mutations, and High, Moderate, and Low influence homozygous mutations. SnpEff annotations were compared between mule deer and previously published white-tailed deer populations (Cars *et al*., 2024).

## Results & Discussion

To contextualize deer diversity and inbreeding patterns, MD and WTD results were compared to previously published estimates for WTD (Cars *et al*., 2024). Genome summary statistics are provided in Table S1. FROH estimates generated with bcftools (model-based) were consistently higher than those obtained using PLINK (rule-based) across WTD populations, although the overall trends were similar (Figure 1; Table S1). Mean FROH (Figure 1) differed among WTD populations, and notably the Island population SPM (x□ = 0.16; 0.0135-0.2389), and the endangered Florida Keys population (x□ = 0.28; 0.0027-0.4752) showed significantly higher estimates than our previous estimate (Cars *et al*., 2024). This is reflective of the bias associated with rule-based ROH methods (Shi et al. 2026; Taylor et al. 2026), as the higher model-based estimates are more consistent with demographic analysis (Kessler et al. 2024) and microsatellite estimates (Villanova et al. 2017). The highlights potential misleading ROH estimates from low-coverage data, and we echo the recommendation for the use of model-based ROH inferences in wild populations.

The mean ROH lengths were similar across populations (Figure 2), with the size distribution consistent with historical bottlenecks rather than contemporary inbreeding (Ceballos et al. 2018). Mule deer had a mean ROH length of 183.8 kb (137.6-239.9 kb) and a mean FROH of 0.07 (0.0057-0.3926). Of note, one MD individual from British Columbia (BTD_BC1) showed significantly higher FROH, which might be indicative of localized inbreeding, though we note ROH values can vary dramatically among individuals within a population (Kardos et al. 2018).

**Figure 2.**
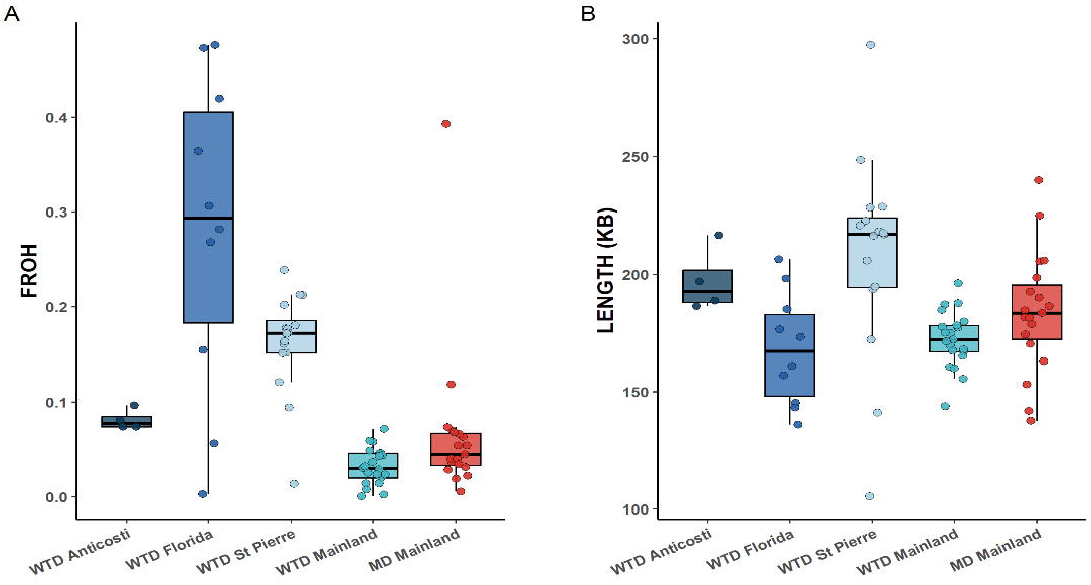
Runs of homozygosity (ROH) estimates in white-tailed deer (*Odocoileus virginianus*) and mule deer (*O. hemionus*) populations: a) is the genome (F) value of inbreeding inferred from ROH; b) is the length distribution of detected ROH.

Mule deer generally had lower genetic load than WTD (Figure 3); here, the long-term low effective population size of MD (Lamb et al. 2021; Kessler et al. 2024) likely has facilitated purging, consisting with other mammals (van der Valk et al. 2019). Though, there were some genetic load outliers in MD, notably individuals from the Pacific Northwest (BTD_BC1, BTD_WA1, MD_OR5, MD_BC8; Figure 1 & 3). The reduced diversity in this region is largely consistent with refugial origins (Latch et al. 2009), but our assessment presents a more nuanced genomic diversity picture compared to previous work (Haynes et al. 2012; Powell et al 2019). The elevated mutation load in the predominantly black-tailed deer region aligns with genome-wide Tajima’s D population estimates (Kessler et al. 2024) and is largely consistent with theoretical predictions (Kardos et al. 2019). This should be monitored going forward.

**Figure 3.**
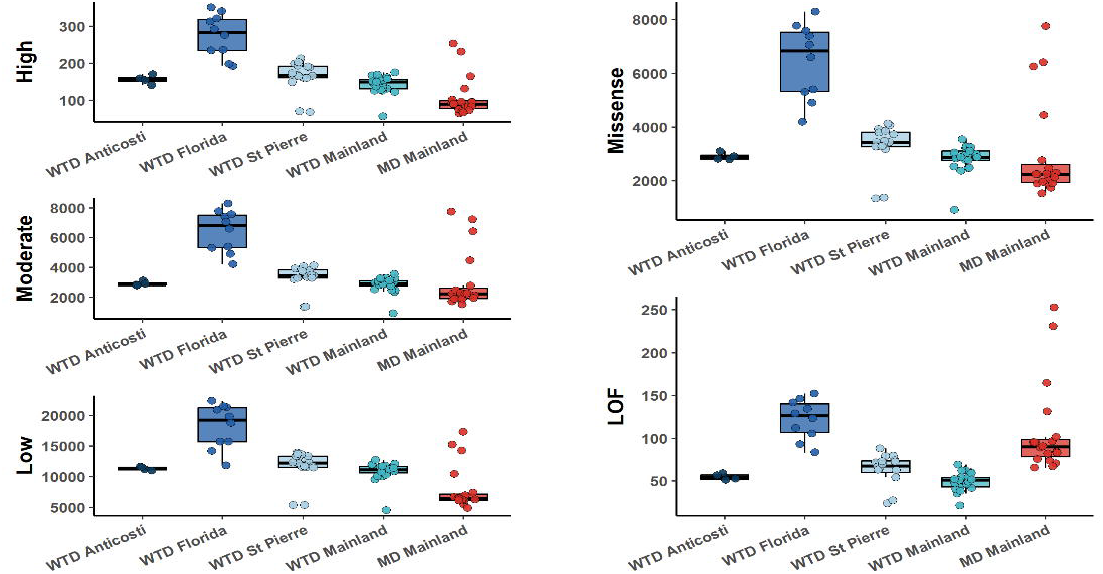
Genetic load estimates in white-tailed deer (*Odocoileus virginianus*) and mule deer (*O. hemionus*) populations: low, moderate and high refer to the predicted impact of the mutation, while missense and loss of function (LOF) are protein impacts.

Interestingly, the number of LOF variants in MD were among the highest observed in our data set (Figure 3). Accumulation of similar defects were thought to contribute to the woolly mammoth extinction (Rogers et al. 2017); here we note the absolute number of LOF mutations are considerably higher in mammoth. By comparison, MD LOF values are close to those reported in Sumatran rhinoceros (Von Seth et al. 2021), but Rhinos have considerably more ROH (Von Seth et al. 2021), so are unable to mask deleterious alleles which is problematic from a conservation standpoint. Given the relatively high effective population sizes of *O. hemionus* (Lamb et al. 2021; Kessler et al. 2024), it is likely these mutations became fixed during a bottleneck, and provided diversity remains as is or increases, should minimize and population-level consequences.

## Acknowledgements

This work was supported by the Natural Sciences and Engineering Research Council of Canada

## Notes

### Competing Interest Statement

The authors have declared no competing interest.

### Summary of Updates

Title change to include common name, grammatical edits to abstract and main text to correct errors and improve flow.

