## Supplementary material for "An updated assessment of the genomic health of *Odocoileus* [deer]": Table S1

|  | **Sample Size** | **Tajima’s *D*** | **Nucleotide  Diversity (π)** | **Mean FROH PLINK** | **Mean FROH Bcftools** | **Mean ROH length (kb)** |
| --- | --- | --- | --- | --- | --- | --- |
| **Anticosti Island** (*O. virginianus*) | 4 | -0.09 | 0.007 | 0.01 | 0.08 | 197.0 |
| **SPM**  (*O. virginianus*) | 16 | 0.54 | 0.005 | 0.02 | 0.16 | 207.9 |
| **Mainland**  (*O. virginianus*) | 21 | -0.39 | 0.006 | 0.01 | 0.03 | 172.3 |
| **Florida Keys** (*O. virginianus*) | 10 | 0.22 | 0.003 | 0.05 | 0.28 | 168.1 |
| **Mainland**  (*O. hemionus*) | 19 | -0.03 | 0.004 | NA | 0.07 | 183.8 |

Table S1. Genome summary statistics for the of white-tailed deer (*Odocoileus virginianus*) and mule deer (*O*. *hemionus*) populations. ROH refers to runs of homozygosity, with F reflecting the inferred inbreeding value.
